# Mutational analysis of N-ethyl-N-nitrosourea (ENU) in the fission yeast *Schizosaccharomyces pombe*

**DOI:** 10.1101/486266

**Authors:** Rafael Hoyos-Manchado, Juan Jiménez, Víctor A. Tallada

## Abstract

Forward genetics has boosted our knowledge on genic function in a multitude of biological models and it has significantly contributed to the understanding of genetic bases of development, ageing and human diseases. With the advent of the next generation sequencing and use of powerful bioinformatic tools, this *traditional* genetic strategy has acquired a new impulse. At present, whole genome sequencing assisted by *in silico* analysis allows the rapid and efficient identification of gene variants that are responsible for a particular phenotype. In this experimental pipeline, it is crucial to start by provoking a large number of random changes in the genome of the model organisms to be screened. A range of chemical mutagens are used to this end because most of them display particular reactivity properties and act differently over DNA. Here we use N-ethyl-N-nitrosourea (ENU) as a mutagen for the first time to our knowledge in the fission yeast *Schizosaccharomyces pombe*. By comparison to the extensively used Ethyl methanesulfonate (EMS) in a phenotype-based study, we conclude that ENU is a very potent and easy-to-use mutagen. Judging from DNA sequence analysis of the identified mutants, ENU induces base changes rather than *indels* and the mutational spectrum in the fission yeast seems similar to that found in mice but different to that described in other single-celled organisms such as budding yeast and *E. coli*. Using ENU in *S. pombe*, we have gathered a collection of 49 auxotrophic mutants including two deleterious alleles of ATIC human ortholog. Defective alleles of this gene are causative of AICA-Ribosiduria, a severe genetic disease. We have also identified 5 aminoglycoside-resistance inactivating mutations in APH genes. All these mutations reported here may be of interest in the metabolism and antibiotic resistance research fields.

## Introduction

The search of a particular phenotype of interest after natural or induced random mutation in biological models, and the consequent identification of the underlying genetic variant, is known as *forward genetics*. This strategy provides an extremely powerful tool to characterise individual gene functions as well as complex genetic pathways interplay, when two or more mutations coexist into the same cell. Human genetic diseases are usually provoked by hypomorphic mutations (partial loss of function) that allow embryo development and birth but become deleterious at later stages. Therefore, by inducing DNA changes into model organisms, genetic bases of human diseases can be very closely reproduced to be studied; even in single-celled organisms such as yeast [1].

A number of physical and chemical mutagens have been used to boost the number of random changes over the natural spontaneous mutation rate. In addition to single base substitutions, most of these agents also provoke small insertions and deletions (generically known as *indels*) [2]. Indels usually lead to frame shifts that generate either truncated or aberrant polypeptides. These provide very valuable information on non-essential gene functions because very often abolish the corresponding protein function entirely or disrupt complexes where they work in, mimicking the full gene deletion phenotype. However, the ideal genetic variants to study essential genes consist of mutations that enable an “on-off’ function switch under given conditions. So called conditional alleles are very often generated by individual amino acid substitutions which “chirurgeonly” destroy postraductional modification sites, unwind binding motives, impair the active centre, etc. without dramatically affecting the overall polypeptide´s structure and/or complexes they are part of. For theses alleles, the switch to destroy function, activity or structure when desired can be turned off simply modulating the culture conditions (temperature, media, presence of chemicals, etc.) [3–5]. These mutations enable elegant genetic studies -otherwise impossible- of essential genes applied to structure-function relationships, biochemical regulation, protein-protein and protein to other partners interactions, etc. Therefore, depending on the aim of the study, it would be desirable to use specific mutagens to enrich the screens for single point substitutions.

Ethyl methanesulfonate (EMS), N-Methyl-N′-nitro-N-nitrosoguanidine (MNNG) and Methyl methanesulfonate (MMS) are within the most common alkylating agents used in yeast and other model organisms as mutation inducing agents [4, 6–8]. Another alkylating agent, N-ethyl-N-nitrosourea (ENU), has also been extensively used for a long time for random mutagenesis screens, especially in mice [9]. However, as far as we know, we are the only ones to use it as a mutagen in fission yeast [3]. As other alkylating counterparts, ENU does not need to be metabolised to alkylate DNA proactively *in vivo* [10–12]. Resulting ethylated nucleotides are mispaired, leading to base pair substitutions after two replication rounds. Importantly, the chemical reactivity described for this molecule remarkably differs from EMS and MMS in the Swain-Scott substrate constant [13, 14]. An especially attractive feature of this molecule is that *Indels* are extremely rare in its mutagenic repertoire [15, 16]; We report here the use of N-ethyl-N-nitrosourea (ENU) as an effective mutagen in the model yeast *Schizosaccharomyces pombe* which may significantly enrich the isolation of new single-base mutant alleles.

## Materials and methods

### Media and growth conditions

Standard fission yeast growth media and molecular biology approaches were used throughout [17]. For qualitative spot tests to assess auxotrophy, every candidate strain was inoculated from a plate in a unique well of a 96-well plate in YES liquid media. After 13 hours, they were diluted 10-fold and plated onto YES and supplemented minimal media lacking adenine, uridine, histidine or leucine in each case. Sporulation agar (SPA) was used for mating and sporulation. Tetrad pulling for segregation analyses was performed in a Singer MSM 400 automated dissection microscope (Singer Instruments).

### Mutagenesis

A reference strain, bearing G418 and Hygromycin B resistance markers integrated into chromosome I and II respectively, was mutagenized using N-ethyl-N-nitrosourea (ENU; Sigma N3385) or ethyl methanesulfonate (EMS; Sigma M0880) as in Winston, 2008 with minor variations [18]. Cells were grown overnight to mid log phase in 500 ml YES medium at 30°C. Cells were concentrated to 2·10^8^ cells/ml and 2 ml aliquots were washed twice with 2 ml of water (centrifuge 5-10 seconds maximum speed), resuspended in 7 ml 0.1 M sodium phosphate buffer (pH 7.0) and split into four aliquots. One of them was treated with 50 ul EMS (0.3 M final) for 1 hour at 30°C, and another one was treated with 60 μl ENU (0.03 M final) for 20 minutes at 30°C. The other two were not treated with mutagens to be used as controls. For ENU treatment and its control, 0.2 ml of each were taken and washed three times with Minimal Medium lacking amino acids and NH_4_ in order to stop mutagenesis. EMS and control aliquots were diluted in 8 ml Na_2_S_2_O_3_ 5% and washed twice in minimal medium lacking amino acids and NH_4_. Cell numbers were scored in Neubauer´s chamber and plated in corresponding dilutions for experiments depicted in Fig 1 (I, II and III).

### Sequencing

Genomic DNA was extracted from selected mutants (Hph^s^, Kan^s^ and Ade/His auxotrophs). Antibiotic resistance cassettes and *ade10* open reading frame were amplified by PCR. Resulting products were sequenced using forward and reverse primers, to ensure total open reading frame coverage in both strands to identify the bases substitutions. Parental strain marker sequencing was used as a control to discard putative previous polymorphism regarding to database sequence.

## Results

### Cellular viability upon ENU vs EMS treatment

In a previous trial for a random mutagenesis study [3] we had roughly adjusted ENU concentration to allow a comparable survival rate to a standard EMS mutagenesis protocol for yeast [18]. Therefore, in the first place, we aimed to finely assess cell viability after ENU treatment in our experimental conditions as compared to well characterised EMS. We cultured our prototroph reference strain (975 background) in yeast extract medium (YES) until log phase (2×10^6^ cells/ml). Cells were harvested and split into four different aliquots of 1×10^8^ cells each (see materials and methods for details and Fig 1) and treated as follows: Aliquot 2 was treated, as in the reference protocol, with 0,3 M EMS for one hour at 30° [18]. Aliquot 4 was treated with 0,03MENU for 20 minutes at 30°. To serve as untreated controls, Aliquots 1 and 3 were processed and washed as aliquots 2 and 4 respectively; except for the lack of the alkylating agent. Afterwards, aliquots 1 and 2 were added into 8 ml of 5% Na_2_S_2_O_3_ to inactivate EMS reactivity and washed immediately with fresh media. We had also found in previous trials that KOH solution, which is used in other systems to inactivate ENU´s reactivity, resulted rather toxic for yeast cells. Thus, aliquots 3 and 4 were washed three times with only minimal medium lacking nitrogen source to avoid cell division before plating. After the last wash, cells were resuspended in fresh media and the number of cells per millilitre was scored in all tubes, considering the average of three independent counts in Neubauer´s chamber as the reference cell number for each tube (Fig 1, Table 1).

**Table 1.**
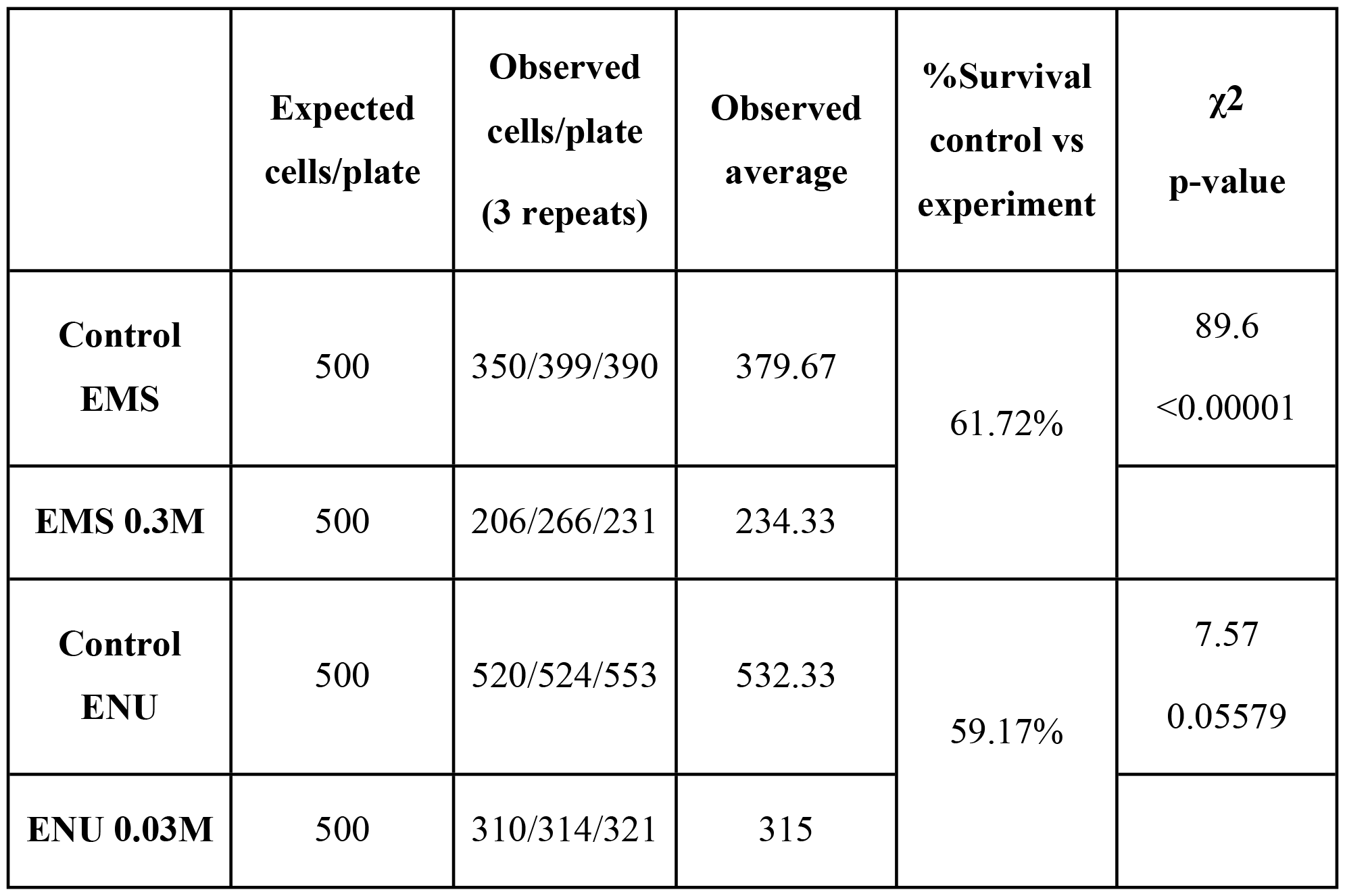
Survival rate after EMS and ENU treatment.

Cell number per ml was estimated for each control and mutagen-containing tubes after the procedure described in materials and methods. Results from three different platings are shown. Respective untreated number of colonies obtained were compared to the expected goal by a chi square test. The result suggests that the treatment used to stop EMS reactivity significantly reduces viability (associated p value<0.00001).

**Fig. 1.**
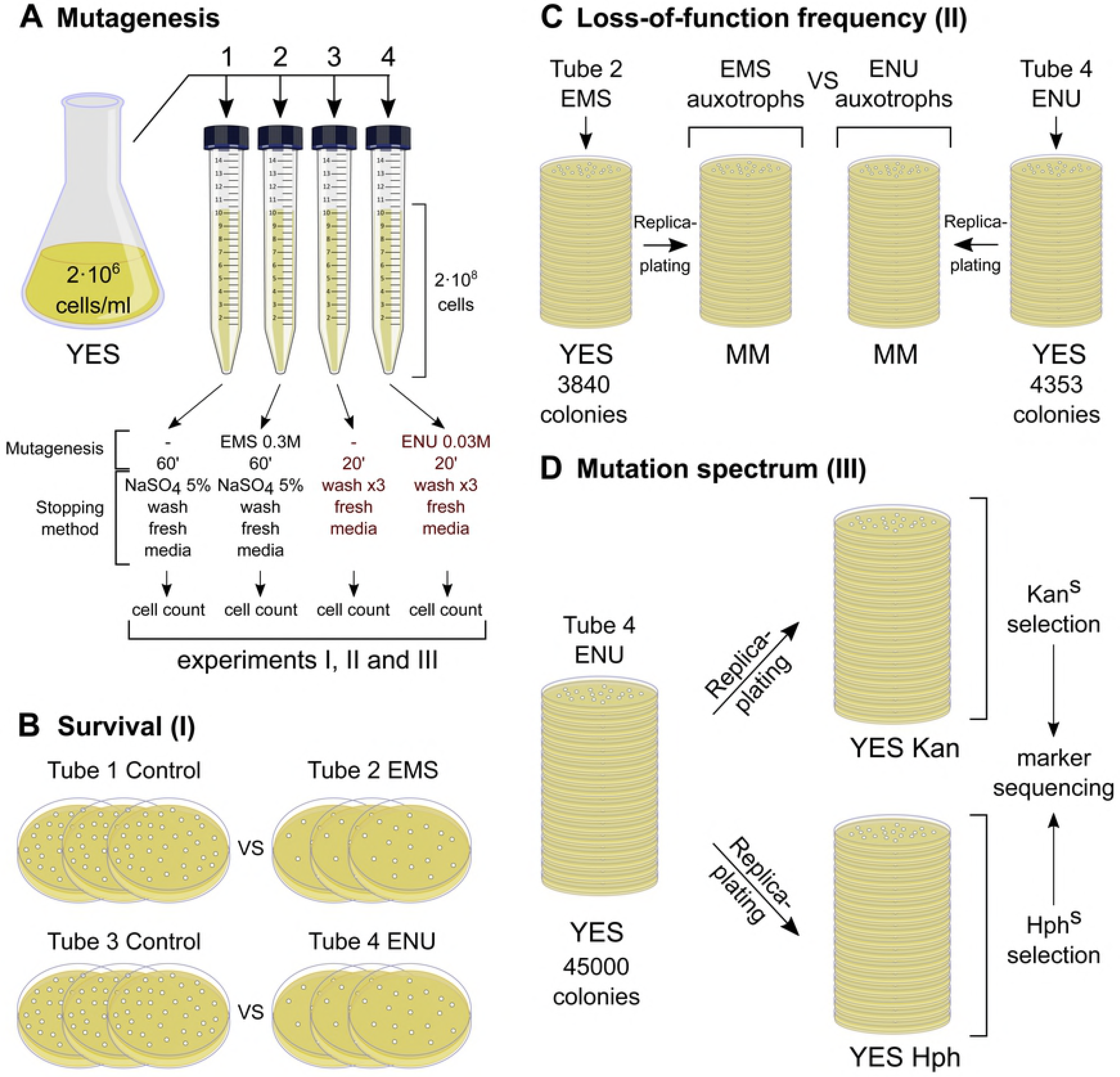
Schematics of the experimental design. (A) Fission yeast cells (975 taxon) bearing KanMX6 and HphMX6 resistance markers were grown in rich media (YES) until early log phase and then split into four tubes. The first and third represent untreated controls while second and fourth were treated with EMS and ENU respectively. (B) Same number of cells, aiming to 500, from each tube in “A” were plated in three technical repeats. Difference in control vs experiment plates was calculated for an estimation of cell survival after exposure to either alkylating agent. (C) Cells exposed to EMS and ENU respectively were grown in solid rich media (YES). Resulting colonies (3840 EMS-treated and 4353 ENU-treated) were replica-plated to synthetic minimal media (MM) without any supplement to identify auxotrophic mutants. (D) Another batch of 45000 ENU-treated colonies were grown in YES media plates as in “C” but replica-plated to YES media containing G418 (indicated as Kan) and Hygromycin B (Hph) antibiotics respectively. Sensitive colonies to either antibiotic were selected to amplify and sequence the resistance marker gene.

To assess survival rate, we plated three technical repetitions for each treatment and controls, aiming to 500 colonies per plate (Fig 1B). As shown in Table 1, both alkylating treatments allow a very similar viability in comparison to their respective untreated control (around 60%); although the exposure time is one third and the molar concentration of ENU is one order of magnitude lower than EMS in this experiment. It is to note as well that the number of viable colonies in aliquot 3 plates (untreated ENU control) are significantly and consistently much closer to the expected goal (500 colonies) than in aliquot 1 (untreated EMS control) (Table 1). This suggests that Na_2_S_2_O_3_ used to inactivate EMS may be also affecting viability itself in spite of such a short exposure time.

### Loss-of-function mutation frequencies

In order to compare the mutagenic potential of ENU and EMS in fission yeast, we screened for auxotrophy-causing mutations as a gene loss-of-function readout over the same number of genomic targets. After EMS and ENU mutagenesis of prototrophic cells explained above (see materials and methods and Fig 1C), we plated them onto YES media. Resulting colonies were counted up and replica-plated onto synthetic minimal media. We then searched for any auxotrophic mutant growing in rich media but unable to proliferate in media with no supplements (EMM). We found 21 such mutants out of 3840 colonies (0,54%) in the case of EMS, and 28 mutants out of 4353 colonies for ENU treatment (0,64%) (Table 2). We further checked if some of these mutants really interrupted specific metabolic pathways and whether any of these could have mutations in more than one metabolic pathway. Mutants were plated onto minimal media lacking just the final product for one of the most common auxotrophic markers used in this yeast: leucine, adenine, uracil and histidine respectively. We found particular auxotrophs for all these metabolites, except for uracil in the case of ENU (Table 2).

**Table 2.** Auxotrophic mutants frequency.

|  | EMS | ENU |
| --- | --- | --- |
| **Leu-** | 1 | 1 |
| **Ura-** | 2 | 0 |
| **Ade-** | 4 | 8* |
| **His-** | 2 | 3* |
| **Other** | 12 | 16 |
| **Total auxotrophs** | 21 | 28 |
| **Colonies analysed** | 3840 | 4353 |
| **Metabolic loss-of-function frequency (%)** | 0.55% | 0.64% |

\* Two of them are both ade- and his-.

Interestingly, two adenine-dependent mutants turned out to be histidine-dependent as well. It is known that both purine and L-histidine synthesis routes share a number of intermediates which make them interdependent [19]. Furthermore, fission yeast *ade9* and *ade10* deletion mutants (adenine dependent) -and their orthologues knock-out in the Baker´s yeast-become histidine dependent as well [19–22]. It is then possible that a single mutation in one of these genes accounts for both auxotrophies. Thus, these could represent ideal candidates for going down to the specific mutation in the DNA to discover new deleterious alleles for both or any of such genes, contributing also to assess the ENU´s mutational spectrum in fission yeast. Therefore, we first crossed both mutants to a wild type strain to check out whether the two metabolic deficiencies segregate together. In both cases, in 16 pulled tetrads, 100% of adenine requiring spores need histidine as well. We then crossed both mutants to each other and did not find any prototroph within the offspring (10 tetrads). These data indicate that there is only one single locus affected and this one is the same in both mutants.

To distinguish among the two putative mutated loci (*ade9* and *ade10*), we checked out genetic linkage to *csi1*, which is just 8 Kb apart from *ade9* (www.pombase.org). Neither mutation showed linkage to *csi1* deletion marker, leaving *ade10* as the only candidate. We therefore sequenced *ade10* coding locus in both mutants. We found a different single base pair substitution in each strain (Table 3), confirming that both mutants are allelic to *ade10* (named *ade10.68* and *ade10.424* respectively) and that the double auxotrophic deficiency is due to a single defective gene rather than two independent mutations in different pathways.

**Table 3.** Mutational spectrum of ENU in fission yeast.

| Marker | Base pair change | Coding nucleotide change<br>(pFA6a numbering) | Amino acid change |
| --- | --- | --- | --- |
| Kan | A-T→G-C | T375C | Val192Ala |
| Kan | A-T→T-A | T626A | Leu209STOP |
| Kan | A-T→G-C | T634C | Cys212Arg |
| Hph | A-T→C-G | A182C | Asp61Ala |
| Hph | G-C→A-T | G293A | Gly98Asp |
| Ade10 | A-T→G-C | T1270C | Ser424Pro |
| Ade10 | A-T→G-C | T203C | Val68Ala |

Base pair substitutions found after ENU treatment are summarized along with the change that abolish function of corresponding protein product.

### Mutational spectrum of ENU in fission yeast

The reference strain used in this study was originally chosen to bear both G418 and Hygromycin B resistance markers integrated into chromosome I and II respectively. To survey for the type of mutations induced by ENU in the fission yeast genome, we selected loss-of-function mutations in those dominant markers after the treatment. To this end, we obtained 45000 colonies in regular YES rich media after ENU treatment described before (tube 4) (Fig 1D). These were replica-plated back to regular YES and YES containing either G418 or Hygromycin B antibiotics (50 mg/ml). Sensitive colonies (three G418^s^ and two Hph^s^) were picked for marker sequencing.

Sequence comparison to wild type control markers identified mutations leading to antibiotic resistance loss (listed in Table 3). Both enzymes conferring G418 and Hygromycin B resistance encode for aminoglycoside phosphotransferases (APH(3')-Ia and APH(4)-Ia types respectively) [23]. Despite a weak sequence identity (12.23%), the spatial protein structure results strikingly similar with two big lobes corresponding roughly to the Amino and Carboxy halves of the protein [24, 25]. Four out of the five mutations target an A-T base pair. All three mutations found in kanMX6 (APH(3')-Ia) lay within a 20 amino acid stretch in the C-terminal domain. T626A base change (KanMX6 numbering) generates a nonsense codon (L209STOP) predicting a 60 amino acid C-terminal truncation. The remaining V192A and C212R amino acid changes fall in close proximity to predicted residues of the aminoglycoside and ATP pockets respectively. In contrast, D61A and G98D changes inactivating HphMX6 (APH(4)-Ia) are located at the N-terminus half, although Glycine 98 is also at the inner face of the nucleotide pocket (https://www.rcsb.org/pdb/protein/P00557?addPDB=3TYK; [24]). These mutations may be of interest in clinic to reveal protein structural changes that inactivate resistance of pathogens to this antibiotic family.

Likewise, the two mutations found in *ade10* open reading frame target two different AT base pairs (transition A-T→G-C) (Table 3). *sp*Ade10 and orthologs encode for a bifunctional enzyme (AICAR formyl transferase/IMP cyclohydrolase) well conserved in prokaryotes and eukaryotes. It uses AICAR (5-Aminoimidazole-4-carboxamide 1-β-D-ribofuranoside, Acadesine, N1-(β-D-Ribofuranosyl) -5-aminoimidazole-4-carboxamide) as the substrate in the last steps of the de novo purine biosynthetic pathway [26, 27]. There are evidences pointing out that an excess of AICAR downregulates histidine synthesis, though the actual biochemical mechanism remains unknown [20]. The deleterious alleles identified in the present study correspond to changes in two very well conserved residues that lead to missense amino acid substitutions with a different expressivity: Val68Ala results a bit leaky while Ser424Pro becomes totally unfunctional. Alignment of 170 orthologs recovered from Ensembl database (https://fungi.ensembl.org/Schizosaccharomycespombe/Gene/ComparaTree/pancompara?g=SPCPB16A4.03c;r=III:947597-949685;t=SPCPB16A4.03c.1) reveals that Valine is the most common residue in position 68 (in 56% of sequences; S1A Fig) and it is present in most eukaryotes within a very well conserved motif (S1B Fig). In position 424, Alanine is the most frequent (57%) residue within all available sequences. However, this residue is present in prokaryotes, mosses and a few plants whilst Serine present in S. pombe is also conserved in nearly all higher eukaryotes including human (31% of database sequences) (S1 Fig). Compound heterozygous deficiency of the human ortholog (ATIC) causes AICA-ribosiduria, a devastating disease characterised by severe neurological defects and congenital blindness. An abnormal accumulation of AICAR and defective purinosome formation is observed in these patients´ fibroblasts [28, 29]. Therefore, the new alleles isolated in this screen could indeed shed light in the study of purine-histidine metabolic pathways crosstalk as well as ATIC-associated ribosiduria.

## Discussion

In order to perform genetic screenings in fission yeast, Ethyl methanesulfonate (EMS), Methyl methanesulfonate (MMS), Nitrosoguanidine (NG), UV radiation and insertional mutagenesis had extensively been used for a long time to cause mutations [4, 30–33]. In high doses, these agents guarantee the generation of a large number of mutants at the cost of a low survival rate. Nevertheless, the spectrum of mutations resulting from UV radiation becomes quite limited and biased [34] and insertional mutagenesis is very useful to disrupt functional elements instead of causing single-base substitutions [33]. On the other hand, DNA alkylating chemicals by no means represent a homogeneous class but differ greatly in their specific bases and atoms attacked and consequently in the potential downstream mutation caused. Alkylation sites in cellular DNA include adenine N^1^, N^3^ and N^7^, guanine N^3^, 0^6^, and N^7^, thymine N^3^, 0^2^ and 0^4^ and cytosine 0^2^ and N^3^. The preference to these sites is largely determined by the nucleophilic selectivity of the agent [35, 36]. It is also well documented that genetic effects of chemical mutagens may vary in different species, strains, sex, age and individuals, even in different cell stages and cell types within the same pluricellular organism [14, 37, 38]. Thus, to enlarge the chemicals shelf that we could use in the fission yeast to induce genetic variations, we decided to implement another alkylating agent, N-ethyl-N-nitrosourea (ENU), since the chemical reactivity of this molecule is qualitatively different from the others mentioned above. Although this agent has been profusely used as a mutagen in mice [39, 40] and to a lesser extent also in other models, including fly [41], worm [42], fish [43], plants [44], bacteria [45] and budding yeast [46]; to our knowledge, this molecule has only been used very few times previously in *S. pombe*, and not as mutagen but as a genotoxin to show sensitivity of a repairing enzyme’s null allele (Atl1) [47, 48]

A very well-established empirical expression of alkylating agents' reactivity is the Swain-Scott substrate constant (*s*). It is a measure of the sensitivity of the alkyl agent to the strength (n) of various nucleophilic reagents with a series of substrates in water solution [13]. Alkylation reactions are then classified in S_N_1-type (low *s*) and S_N_2-type (high *s*). Most of alkylating compounds used as mutagens range from *s*=0,26 to *s*=0,86. Both EMS and ENU act by transferring the ethyl group into nucleotides. However, there is a remarkable difference in their reactivity: whilst EMS act by a mixed S_N_1/S_N_2-type reaction and shows a relatively high Swain-Scott constant (*s*=0.67), ENU only acts by a S_N_1-type reaction with a much lower *s* constant (*s*=0.26) [49]. This translates into EMS attacking mostly high nucleophilicity ring N sites (mostly the N^7^ position of guanine) and O-atoms up to certain extent. Therefore, EMS mutagenesis is biased towards G/C to A/T transitions which often leads to the generation of stop codons [50]. It has also been reported that EMS can generate up to 13% of deletions and other chromosome rearrangements [51], some double strand breaks and attacking also proteins to a considerable extent [52]. On the other hand, ENU acts preferentially towards low nucleophilicity O-atoms in DNA (mainly O^2^ and O^4^ in thymine and O^6^-deoxyguanosine) [14, 53]. In order to compare the action of two chemicals it is important to consider the trade-off between genotoxic power and population survival. For this comparison, we worked out the conditions allowing almost identical survival rate treating with EMS and ENU respectively. This implied that the molar concentration of ENU had to be adjusted at one order of magnitude lower than EMS and time of exposure reduced to a third. This suggests that ENU results a rather more genotoxic agent than EMS in fission yeast. Both the higher killing power and mutagenic potential over EMS have been previously observed in other systems [14, 54].

In the present gene-based study we have also surveyed for ENU´s mutational spectrum. A number systematic and gene/phenotype-based studies reports on the range of ENU induced substitutions have been previously carried out with different outcomes. In *E. coli*, *S. cerevisiae* and *C. elegans* over 70% of changes corresponded to G-C to AT transitions while almost the remaining 30% were A-T to T-A transversions [46, 55, 56]. However other studies in mouse (compiled in [15, 57, 58]) and a systematic study in toxoplasma, revealed opposite proportions: more than 75% of changes affected A-T pairs [59]. Furthermore, in *Drosophila* germ cells selectivity became one or another depending on the pre-or-post meiotic stage [41, 60, 61]. This could be explained under a generally accepted concept suggesting that 0^6^-alkylguanine and 0^4^-alkylthymine have promutagenic potential towards transition mutations (GC→AT and AT→GC, respectively) [62, 63] and 0^2^-alkylthymine to AT→TA transversions [64]. In this scenario, the protective and repairing enzymes background, cell cycle stage and mutagen doses might be important bias factors [48]. In this study all mutations identified correspond to single base pair substitutions (Two in *ade10*, two in APH(4)-Ia and three in APH(3')-Ia). Six out of seven mutations inactivating each phosphotransferase conferring antibiotic resistance or the *ade10* gene, are transitions or transversions from a native A-T pair (Table 3). In spite of the limited number of mutations analysed, all observed changes in other models are represented in this study but the preference in fission yeast seems to be in line with models such as mouse and toxoplasma rather than *E. coli* or *S. cerevisiae*. As mentioned above, this could be related with the specificity and efficiency of alkyltransferase-like protective proteins in *S. pombe* such as Atl1 against 0^6^-alkylguanine adducts [47, 48, 65].

Nevertheless, in contrast to other alkylating agents, indels imputed to ENU are extremely rare or absent in literature in any model, [16, 46]. This implies that loss-of-function mutations generated by ENU are caused by single base substitutions rather than by frame shifts that usually result more deleterious. Furthermore, in phenotype and gene-driven screens it is also evident that, within the substitutions category, missense mutations are much more likely to occur than protein-truncating nonsense mutations [15]. This sets a remarkable difference between ENU and other alkylating counterparts to be used to complement each other to enlarge the changes spectrum in mutational studies. It has also been shown by whole genome sequencing studies that ENU induced mutations are randomly distributed over a whole genome [59]. In our loss-of-function mutations screens, on the one hand, over about 93 potential target genes (listed in fission yeast phenotype ontology database as auxotrophy-causing, PomBase FYPO:0000128) we found 28 auxotrophic mutants in 4353 colonies. On the other hand, over two target dominant markers, we identified 5 mutants in 45000 colonies. Therefore, these experimental conditions would give a rough estimation between 14520 and 18000 colonies to be screened to find a knock-out hit in a given average gene. Slight variations are to be considered by factors such as G-C content or gene´s size.

Taking all this together, we propose that this molecule can be very efficiently used in mutagenesis studies in *Schizosaccharomyces pombe* (and very likely in other prominent yeast and fungi models such as *Candida albicans*, *Ustilago maydis*, etc.), either as an alternative or as a complement to its sulfonate and nitroso counterparts. In addition to finding new valuable conditional mutations in forward genetics screens, this could contribute in directed evolution experiments, suppression studies etc. where subtle changes are desired. As a matter of fact, we have gathered in this study a collection of 49 auxotrophic mutants and identified 3 and 2 base substitutions which inactivate respectively APH(3')-Ia and APH(4)-Ia phosphotransferases genes. In the case of auxotrophs, it becomes somehow surprising that among a limited (and nonsaturating) number of mutations generated here, two of them fall down within the same locus (*ade10*). However in a meta-study of mutagenic spectrum of ENU in mouse, Barbaric et al found that genes with higher G-C content become more likely to be mutated by ENU and showed that A-T pairs flanked by C or G bases were more prone to mutation [15]. Notably, *ade10* open reading frame contains the highest G-C percentage (46%) of the whole adenine pathway and significantly higher than fission yeast protein-coding gene average G-C content (37.08%) [66]. This might suggest that gene-wise, ENU-induced changes frequency could be biased by G-C content also in fission yeast. Although not in the scope of this work, all mutants reported here might be of interest in the study of metabolic diseases and antibiotic resistance research fields.

## Acknowledgments

We thank Víctor Carranco and Anabel Lopez for excellent technical support, Alejandro Rubio for sequence alignment assistance and Andrés Garzón for critical reading of the manuscript.

## Supporting information

**S1 Fig. Evolutionary conservation of ADE10/ATIC gene.** A) Frequencies of residues found in position 68 and 424 (*S. pombe* numbering) within all ortholog sequences available in Ensembl database. B) Alignment of respective surrounding sequence stretches around residues 68 and 424. Representative prokaryotic and eukaryotic model organisms are included. Valine and Serine present in *S. pombe* are tightly conserved in other yeasts and animals including human.

## References

1. Tenreiro S, Franssens V, Winderickx J, Outeiro TF. Yeast models of Parkinson’s disease-associated molecular pathologies. Current Opinion in Genetics and Development. 2017;44:74–83. doi: 10.1016/j.gde.2017.01.013. PubMed PMID: 28232272.

2. Henry IM, Nagalakshmi U, Lieberman MC, Ngo KJ, Krasileva KV, Vasquez-Gross H, et al. Efficient Genome-Wide Detection and Cataloging of EMS-Induced Mutations Using Exome Capture and Next-Generation Sequencing. The Plant Cell. 2014;26(4):1382–97. doi: 10.1105/tpc.113.121590. PubMed PMID: 24728647; PubMed Central PMCID: PMC4036560.

3. Hoyos-Manchado R, Reyes-Martin F, Rallis C, Gamero-Estevez E, Rodriguez-Gomez P, Quintero-Blanco J, et al. RNA metabolism is the primary target of formamide in vivo. Sci Rep. 2017;7(1):15895. doi: 10.1038/s41598-017-16291-8. PubMed PMID: 29162938; PubMed Central PMCID: PMC5698326.

4. Bonatti S, Simili M, Abbondandolo A. Isolation of temperature-sensitive mutants of Schizosaccharomyces pombe. J Bacteriol. 1972;109(2):484–91. PubMed PMID: 4110142; PubMed Central PMCID: PMC285166.

5. Willis JH, Munro E, Lyczak R, Bowerman B. Conditional dominant mutations in the Caenorhabditis elegans gene act-2 identify cytoplasmic and muscle roles for a redundant actin isoform. Molecular biology of the cell. 2006;17(3):1051–64. doi: 10.1091/mbc.e05-09-0886. PubMed PMID: 16407404; PubMed Central PMCID: PMC1382297.

6. Segal E, Munz P, Leupold U. Characterization of chemically induced mutations in the ad-1 locus of Schizosaccharomyces pombe. Mutation Research - Fundamental and Molecular Mechanisms of Mutagenesis. 1973;18(1):15–24. doi: 10.1016/0027-5107(73)90017-1.

7. Spencer JF, Spencer DM. Mutagenesis in yeast. Methods in molecular biology. 1996;53:17–38. doi: 10.1385/0-89603-319-8:17. PubMed PMID: 8924979.

8. Sabatinos SA, Forsburg SL. Chapter 32 - Molecular Genetics of Schizosaccharomyces pombe. Methods in Enzymology. 470: Academic Press; 2010. p. 759–95.

9. Russell WL, Kelly EM, Hunsicker PR, Bangham JW, Maddux SC, Phipps EL. Specific-locus test shows ethylnitrosourea to be the most potent mutagen in the mouse. Proceedings of the National Academy of Sciences. 1979;76(11):5818–9. doi: 10.1073/pnas.76.11.5818. PubMed PMID: 293686; PubMed Central PMCID: PMC411742.

10. Haglund J, Van Dongen W, Lemière F, Esmans EL. Analysis of DNA-phosphate adducts in vitro using miniaturized LC-ESI-MS/MS and column switching: Phosphotriesters and alkyl cobalamins. Journal of the American Society for Mass Spectrometry. 2004;15(4):593–606. doi: 10.1016/j.jasms.2003.12.012. PubMed PMID: 15047064.

11. Wani AA, Gibson-D’Ambrosio RE, D’Ambrosio SM. Quantitation of O6-ethyldeoxyguanosine in ENU alkylated DNA by polyclonal and monoclonal antibodies. Carcinogenesis. 1984;5(9):1145–50. Epub 1984/09/01. PubMed PMID: 6467504.

12. Scalera SE, Ward OG. A quantitative study of ethyl methanesulfonate-induced alkylation of Vicia faba DNA. Mutation Research/Fundamental and Molecular Mechanisms of Mutagenesis. 1971;12(1):71–9. doi: 10.1016/0027-5107(71)90074-1.

13. Swain CG, Scott CB. Quantitative Correlation of Relative Rates. Comparison of Hydroxide Ion with Other Nucleophilic Reagents toward Alkyl Halides, Esters, Epoxides and Acyl Halides1. Journal of the American Chemical Society. 1953;75(1):141–7. doi: 10.1021/ja01097a041.

14. Vogel EW, Natarajan AT. The Relation between Reaction Kinetics and Mutagenic Action of Monofunctional Alkylating Agents in Higher Eukaryotic Systems: Interspecies Comparisons. In: Serres FJ, Hollaender A, editors. Chemical Mutagens. 7. New York: Plenum Press; 1982. p. 295–336.

15. Barbaric I, Wells S, Russ A, Dear TN. Spectrum of ENU-induced mutations in phenotype-driven and gene-driven screens in the mouse. Environmental and molecular mutagenesis. 2007;48(2):124–42. doi: 10.1002/em.20286. PubMed PMID: 17295309.

16. Takahasi KR, Sakuraba Y, Gondo Y. Mutational pattern and frequency of induced nucleotide changes in mouse ENU mutagenesis. BMC Molecular Biology. 2007;8:1–10. doi: 10.1186/1471-2199-8-52. PubMed PMID: 17584492; PubMed Central PMCID: PMC1914352.

17. Moreno S, Klar A, Nurse P. Molecular genetic analysis of fission yeast Schizosaccharomyces pombe. Methods in enzymology. 1991;194:795–823. PubMed PMID: 2005825.

18. Winston F. EMS and UV mutagenesis in yeast. Current Protocols in Molecular Biology. 2008;Chapter 13:1–5. doi: 10.1002/0471142727.mb1303bs82. PubMed PMID: 18425760.

19. Tibbetts AS, Appling DR. Characterization of Two 5-Aminoimidazole-4-carboxamide Ribonucleotide Transformylase/Inosine Monophosphate Cyclohydrolase Isozymes from Saccharomyces cerevisiae. Journal of Biological Chemistry. 2000;275(27):20920–7. doi: 10.1074/jbc.M909851199. PubMed PMID: 10877846.

20. Rébora K, Laloo B, Daignan-Fornier B. Revisiting Purine-Histidine Cross Pathway Regulation in Saccharomyces cerevisiae. A Central Role for a Small Molecule. 2005;170(1):61–70. doi: 10.1534/genetics.104.039396. PubMed PMID: 15744050; PubMed Central PMCID: PMC1449729.

21. Jones EW, Fink GR. Regulation of Amino Acid and Nucleotide Biosynthesis in Yeast. In: Strathern JN, Jones EW, Broach JR, editors. The Molecular Biology of the Yeast Saccharomyces: Metabolism and Gene Expression.: Cold Spring Harbor; 1982.

22. Whitehead E, Nagy M, Heslot H. Interactions entre la biosynlhèse des purines nucléotides et celle de l'histidine chez le Schizosaccharomyces pombe. Comptes rendus hebdomadaires des seances de l'Academie des sciences Serie D: Sciences naturelles. 1966;263:819–21.

23. Wright GD, Thompson PR. Aminoglycoside phosphotransferases: proteins, structure, and mechanism. Front Biosci. 1999;4:D9–21. doi: 10.2741/Wright. PubMed PMID: 9872733.

24. Stogios PJ, Shakya T, Evdokimova E, Savchenko A, Wright GD. Structure and function of APH(4)-Ia, a hygromycin B resistance enzyme. Journal of Biological Chemistry. 2011;286(3):1966–75. doi: 10.1074/jbc.M110.194266. PubMed PMID: 21084294; PubMed Central PMCID: PMC3023493.

25. Stogios PJ, Spanogiannopoulos P, Evdokimova E, Egorova O, Shakya T, Todorovic N, et al. Structure-guided optimization of protein kinase inhibitors reverses aminoglycoside antibiotic resistance. The Biochemical journal. 2013;454(2):191–200. doi: 10.1042/BJ20130317. PubMed PMID: 23758273; PubMed Central PMCID: PMC3743924.

26. Rayl EA, Moroson BA, Beardsley GP. The Human purH Gene Product, 5-Aminoimidazole-4-carboxamide Ribonucleotide Formyltransferase/IMP Cyclohydrolase. Cloning, sequencing, expression, purification, kinetic analysis, and domain mapping. Journal of Biological Chemistry. 1996;271(4):2225–33. doi: 10.1074/jbc.271.4.2225. PubMed PMID: 8567683.

27. Richter R, Heslot H. Genetic and functional analysis of the complex locus ade10 in Schizosaccharomyces pombe. Current Genetics. 1982;5(3):233–44. doi: 10.1007/BF00391812. PubMed PMID: 24186301.

28. Baresova V, Skopova V, Sikora J, Patterson D, Sovova J, Zikanova M, et al. Mutations of ATIC and ADSL affect purinosome assembly in cultured skin fibroblasts from patients with AICA-ribosiduria and ADSL deficiency. Hum Mol Genet. 2012;21(7):1534–43. doi: 10.1093/hmg/ddr591. PubMed PMID: 22180458.

29. Marie S, Heron B, Bitoun P, Timmerman T, Van den Berghe G, Vincent M-F. AICA-Ribosiduria: A Novel, Neurologically Devastating Inborn Error of Purine Biosynthesis Caused by Mutation of ATIC. The American Journal of Human Genetics. 2004;74(6):1276–81. doi: 10.1086/421475. PubMed PMID: 15114530; PubMed Central PMCID: PMC1182092.

30. Berry CH, Ibrahim MA, Coddington A. Characterisation of ribosomes from drug resistant strains of Schizosaccharomyces pombe in a poly U directed cell free protein synthesising system. Mol Gen Genet. 1978;167(2):217–25. doi: 10.1007/BF00266915. PubMed PMID: 732808.

31. Loprieno N, Guglielminetti R, Bonatti S, Abbondandolo A. Evaluation of the genetic alterations induced by chemical mutagens in Schizosaccharomyces pombe. Mutation Research/Fundamental and Molecular Mechanisms of Mutagenesis. 1969;8(1):65–71. doi: http://dx.doi.org/10.1016/0027-5107(69)90141-9. PubMed PMID: 5796945.

32. Potashkin J, Li R, Frendewey D. Pre-mRNA splicing mutants of Schizosaccharomyces pombe. The EMBO Journal. 1989;8(2):551–9. PubMed PMID: 400840; PubMed Central PMCID: PMC400840.

33. Chua G, Taricani L, Stangle W, Young PG. Insertional mutagenesis based on illegitimate recombination in Schizosaccharomyces pombe. Nucleic Acids Res. 2000;28(11):E53. PubMed PMID: 10871352; PubMed Central PMCID: PMC102638.

34. Pfeifer GP, You YH, Besaratinia A. Mutations induced by ultraviolet light. Mutation Research - Fundamental and Molecular Mechanisms of Mutagenesis. 2005;571(1-2):19–31. doi: 10.1016/j.mrfmmm.2004.06.057. PubMed PMID: 15748635.

35. Beranek DT. Distribution of methyl and ethyl adducts following alkylation with monofunctional alkylating agents. Mutation Research/Fundamental and Molecular Mechanisms of Mutagenesis. 1990;231(1):11–30. doi: https://doi.org/10.1016/0027-5107(90)90173-2. PubMed PMID: 2195323.

36. Vogel EW, Natarajan AT. DNA damage and repair in somatic and germ cells in vivo. Mutation Research - Fundamental and Molecular Mechanisms of Mutagenesis. 1995;330(1-2):183–208. doi: 10.1016/0027-5107(95)00040-P. PubMed PMID: 7623865.

37. Gibson-D'Ambrosio RE, Leong Y, D'Ambrosio SM. DNA Repair following Ultraviolet and N-Ethyl-N-nitrosourea Treatment of Cells Cultured from Human Fetal Brain, Intestine, Kidney, Liver, and Skin. Cancer Research. 1983;43(12 Part 1):5846–50.

38. Slikker W, Andersen ME, Bogdanffy MS, Bus JS, Cohen SD, Conolly RB, et al. Dose-dependent transitions in mechanisms of toxicity. Toxicology and Applied Pharmacology. 2004;201(3):203–25. doi: https://doi.org/10.1016/j.taap.2004.06.019. PubMed PMID: 15582645.

39. Masumura K, Matsui M, Katoh M, Horiya N, Ueda O, Tanabe H, et al. Spectra of gpt mutations in ethylnitrosourea-treated and untreated transgenic mice. Environmental and molecular mutagenesis. 1999;34(1): 1–8. doi: doi: 10.1002/(SICI)1098-2280(1999)34:1<1::AID-EM1>3.0.C0;2-P. PubMed PMID: 10462717.

40. Salinger AP, Justice MJ. Mouse Mutagenesis Using N-Ethyl-N-Nitrosourea (ENU). Cold Spring Harbor Protocols. 2008;2008(4):pdb.prot4985. doi: 10.1101/pdb.prot4985. PubMed PMID: 21356809.

41. Pastink A, Vreeken C, Nivard MJ, Searles LL, Vogel EW. Sequence analysis of N-ethyl-N-nitrosourea-induced vermilion mutations in Drosophila melanogaster. Genetics. 1989;123(1):123–9. PubMed PMID: 2572507; PubMed Central PMCID: PMC1203775.

42. De Stasio EA, Dorman S. Optimization of ENU mutagenesis of Caenorhabditis elegans. Mutation Research/Genetic Toxicology and Environmental Mutagenesis. 2001;495(1):81–8. doi: https://doi.org/10.1016/S1383-5718(01)00198-X. PubMed PMID: 11448645.

43. de Bruijn E, Cuppen E, Feitsma H. Highly Efficient ENU Mutagenesis in Zebrafish. In: Lieschke GJ, Oates AC, Kawakami K, editors. Zebrafish: Methods and Protocols. Totowa, NJ: Humana Press; 2009. p. 3–12.

44. Gichner T. Differential genotoxicity of ethyl methanesulphonate, N-ethyl-N-nitrosourea and maleic hydrazide in tobacco seedlings based on data of the Comet assay and two recombination assays. Mutation Research/Genetic Toxicology and Environmental Mutagenesis. 2003;538(1):171–9. doi: https://doi.org/10.1016/S1383-5718(03)00117-7. PubMed PMID: 12834766.

45. Hince TA, Neale S. A comparison of the mutagenic action of the methyl and ethyl derivatives of nitrosamides and nitrosamidines on Escherichia coli. Mutation Research/Fundamental and Molecular Mechanisms of Mutagenesis. 1974;24(3):383–7. doi: https://doi.org/10.1016/0027-5107(74)90183-3. PubMed PMID: 4606103.

46. Lee GSF, Blonsky KS, Van On DL, Savage EA, Morgan AR, von Borstel RC. Base alterations in yeast induced by alkylating agents with differing Swain-Scott substrate constants. Journal of Molecular Biology. 1992;223(3):617–26. doi: https://doi.org/10.1016/0022-2836(92)90978-S.

47. Latypov VF, Tubbs JL, Watson AJ, Marriott AS, McGown G, Thorncroft M, et al. Atl1 regulates choice between global genome and transcription-coupled repair of O(6)-alkylguanines. Molecular cell. 2012;47(1):50–60. Epub 05/31. doi: 10.1016/j.molcel.2012.04.028. PubMed PMID: 22658721.

48. Pearson SJ, Wharton S, Watson AJ, Begum G, Butt A, Glynn N, et al. A novel DNA damage recognition protein in Schizosaccharomyces pombe. Nucleic acids research. 2006;34(8):2347–54. doi: 10.1093/nar/gkl270. PubMed PMID: 16679453; PubMed Central PMCID: PMC1458281.

49. Boffa LC, Bolognesi C, Mariani MR. Specific targets of alkylating agents in nuclear proteins of cultured hepatocytes. Mutation research. 1987;190(2):119–23. doi: 10.1016/0165-7992(87)90042-X. PubMed PMID: 3821770.

50. Flibotte S, Edgley ML, Chaudhry I, Taylor J, Neil SE, Rogula A, et al. Whole-Genome Profiling of Mutagenesis in Caenorhabditis elegans. Genetics. 2010;185(2):431–41. doi: 10.1534/genetics.110.116616. PubMed PMID: 20439774; PubMed Central PMCID: PMC2881127.

51. Anderson P. Mutagenesis. Methods in cell biology. 1995;48:31–58. Epub 1995/01/01. PubMed PMID: 8531732.

52. Sega GA. A review of the genetic effects of ethyl methanesulfonate. Mutation Research/Reviews in Genetic Toxicology. 1984;134(2-3):113–42. doi: https://doi.org/10.1016/0165-1110(84)90007-1. PubMed PMID: 6390190.

53. Loveless A. Possible relevance of O-6 alkylation of deoxyguanosine to the mutagenicity and carcinogenicity of nitrosamines and nitrosamides. Nature. 1969;223(5202):206–7. Epub 1969/07/12. PubMed PMID: 5791738.

54. Natarajan AT, Simons JWIM, Vogel EW, van Zeeland AA. Relationship between cell killing, chromosomal aberrations, sister-chromatid exchanges and point mutations induced by monofunctional alkylating agents in Chinese hamster cells a correlation with different ethylation products in DNA. Mutation Research - Fundamental and Molecular Mechanisms of Mutagenesis. 1984;128(1):31–40. doi: 10.1016/0027-5107(84)90044-7. PubMed PMID: 6472304.

55. Burns PA, Gordon AJE, Kunsmann K, Glickman BW. Influence of Neighboring Base Sequence on the Distribution and Repair of N-Ethyl-N-nitrosourea-induced Lesions in Escherichia coli. Cancer Research. 1988;48(16):4455–8.

56. Sarin S, Bertrand V, Bigelow H, Boyanov A, Doitsidou M, Poole RJ, et al. Analysis of multiple ethyl methanesulfonate-mutagenized Caenorhabditis elegans strains by whole-genome sequencing. Genetics. 2010;185(2):417–30. doi: 10.1534/genetics.110.116319. PubMed PMID: 20439776; PubMed Central PMCID: PMC2881126.

57. Arnold CN, Barnes MJ, Berger M, Blasius AL, Brandl K, Croker B, et al. ENU-induced phenovariance in mice: inferences from 587 mutations. BMC Res Notes. 2012;5:577. doi: 10.1186/1756-0500-5-577. PubMed PMID: 23095377; PubMed Central PMCID: PMC3532239.

58. Noveroske JK, Weber JS, Justice MJ. The mutagenic action of N-ethyl-N-nitrosourea in the mouse. Mammalian Genome. 2000;11(7):478–83. doi: 10.1007/s003350010093. PubMed PMID: 10886009.

59. Farrell A, Coleman BI, Benenati B, Brown KM, Blader IJ, Marth GT, et al. Whole genome profiling of spontaneous and chemically induced mutations in Toxoplasma gondii. BMC Genomics. 2014;15:1–15. doi: 10.1186/1471-2164-15-354. PubMed PMID: 24885922; PubMed Central PMCID: PMC4035079.

60. Fossett NG, Arbour-Reily P, Kilroy G, McDaniel M, Mahmoud J, Tucker AB, et al. Analysis of ENU-induced mutations at the Adh locus in Drosophila melanogaster. Mutation Research - Fundamental and Molecular Mechanisms of Mutagenesis. 1990;231(1):73–85. doi: 10.1016/0027-5107(90)90178-7. PubMed PMID: 2114535.

61. Tosal L, Comendador MA, Sierra LM. N-ethyl-N-nitrosourea predominantly induces mutations at AT base pairs in pre-meiotic germ cells of Drosophila males. Mutagenesis. >1998;13(4):375–80. Epub 1998/08/26. PubMed PMID: 9717174.

62. Justice MJ, Noveroske JK, Weber JS, Zheng B, Bradley A. Mouse ENU Mutagenesis. Human Molecular Genetics. 1999;8(10):1955–63. doi: 10.1093/hmg/8.10.1955. PubMed PMID: 10469849.

63. Preston BD, Singer B, Loeb LA. Comparison of the relative mutagenicities of O-alkylthymines site-specifically incorporated into phi X174 DNA. The Journal of biological chemistry. 1987;262(28):13821–7. PubMed PMID: 2958453.

64. Grevatt PC, Solomon JJ, Bhanot OS. In Vitro Mispairing Specificity of O2-Ethylthymidinet. Biochemistry. 1992;31(17):4181–8. doi: 10.1021/bi00132a005.

65. Bronstein SM, Skopek TR, Swenberg JA. Efficient Repair of O6-Ethylguanine, but not O4-Ethylthymine or O2-Ethylthymine, Is Dependent upon O6-Alkylguanine-DNA Alkyltransferase and Nucleotide Excision Repair Activities in human cells. Cancer Research. 1992;52:2008 LP - 11.

66. Lock A, Rutherford K, Harris MA, Hayles J, Oliver SG, Bähler J, et al. PomBase 2018: user-driven reimplementation of the fission yeast database provides rapid and intuitive access to diverse, interconnected information. Nucleic Acids Research. 2018:1–7. doi: 10.1093/nar/gky961. PubMed PMID: 30321395.

